# Pseudoassembly of k-mers

**DOI:** 10.1101/2025.05.11.653354

**Authors:** Delaney K. Sullivan, Mayuko Boffelli, Lior Pachter

## Abstract

We introduce a *pseudoassembly* approach to identifying variation in sets of genomic sequences via colored de Bruijn graphs. Our pseudoassembly method is implemented in a program called klue that assembles k-mers into sequences compatible with a variant-aware extension of pseudoalignment. We show that this approach can be used to identify cell-type specific *de novo* variants from single-cell RNA-seq in a mouse melanoma model.

## Introduction

The de Bruijn graph has served as an effective data structure for pseudoalignment (Bray et al. 2016; Almodaresi, Zakeri, and Patro 2021) by compressing the information embedded in k-mers present in target sequences that read sequences will be compared to. In this application, each k-mer, indexed within a colored de Bruijn graph (Alekseyev and Pevzner 2007; Iqbal et al. 2012), originates from one or more target sequences, typically extracted from an annotation of a reference genome. An extension of this framework introduced in (Sullivan et al. 2025) uses unannotated k-mers, called distinguishing flanking k-mers, which may be extracted from unannotated sequences, e.g. directly from a genome. Such k-mers have proved valuable in improving the accuracy of pseudoalignment.

The (Sullivan et al. 2025) idea can be generalized and systematized: Rather than relying on known annotations, target sequences can be constructed *de novo* via structural searching of colored de Bruijn graphs built from multiple types of sequences. We refer to this approach as pseudoassembly, in the spirit of the term pseudoalignment. Unlike full assembly, which reconstructs entire transcripts or genomes, pseudoassembly focuses on only indexing k-mers that are relevant for downstream read assignment. This targeted construction avoids the complexities of full assembly while retaining the ability to detect biologically meaningful sequence variation.

Pseudoassembly is particularly valuable for studying genetic variation. Because k-mers can represent variant sequences, they are well-suited for identifying sample-specific or condition-specific differences in sequence content. One common strategy for analyzing known variants involves tiling k-mer windows around known polymorphisms, such as SNPs, structural variants, or splice junctions, and indexing them for downstream analysis. This approach depends on prior knowledge of variant locations. In contrast, pseudoassembly proceeds *de novo* from raw sequences in FASTQ or FASTA format. For instance, the pseudoassembly can facilitate the extraction of k-mers that are uniquely found in an “experimental” sample, e.g. sequences absent from control datasets or the reference genome). This is particularly relevant in settings such as cancer, where mutations are highly variable in both structure and genomic location. The flexibility and reference-agnostic nature of k-mers make them ideal for capturing such unstructured variation.

Several prior tools have utilized k-mers to directly probe differences between sequence sets. For example, DE-kupl focuses on differential k-mer abundance between conditions and has demonstrated high sensitivity in detecting novel biological variation (Audoux et al. 2017) and differential k-mers were used by (Rahman et al. 2018) for association mapping. Another more recent method, SPLASH, uses “anchor” and “target” k-mers, where a sequence anchor may be associated with multiple different targets (each representing a variant), to quantify variation (Chaung et al. 2023).

Building on these ideas, we developed klue (<u>k</u>-mer based <u>l</u>ocal <u>u</u>niqueness <u>e</u>xploration), a general-purpose tool based on pseudoassembly for discovering, organizing, and extracting informative k-mers. klue accepts both FASTQ and FASTA files as input and can be applied to RNA-seq data, DNA-seq data, or sequence assemblies.

## Results

### Overview

The klue method uses k-mers that are extracted in the form of longer contiguous sequences (contigs). Following contig extraction (where the contigs represent variant-specific k-mers), klue proceeds to pseudoassembly. This involves mapping the contigs of interest—those extracted by klue—to known reference sequences. For example, if the input k-mer size was 31, smaller k-mer matches (e.g. 29-mers) might be used to find a partial match between a contig and annotated transcript sequences. When a contig is mapped to a target sequence, it inherits the color of that target in a de Bruijn graph. However, to distinguish a contig as a variant-derived sequence, the color is modified into a uniquely shaded version. Each distinct contig mapped to a target gets its own unique shade. If a contig maps to multiple target sequences, it is assigned a shade for each.

This process gives rise to an ornamental de Bruijn graph (Figure 1), a concept that was introduced in earlier work and implemented in the tool Ornaments (Adduri and Kim 2024). In this setup, pseudoalignment proceeds as usual via set intersection of k-mer colors. However, a set union is performed for the shades that are encountered in a read. The resulting equivalence class will contain both the parent colors and the shades.

**Figure 1:**
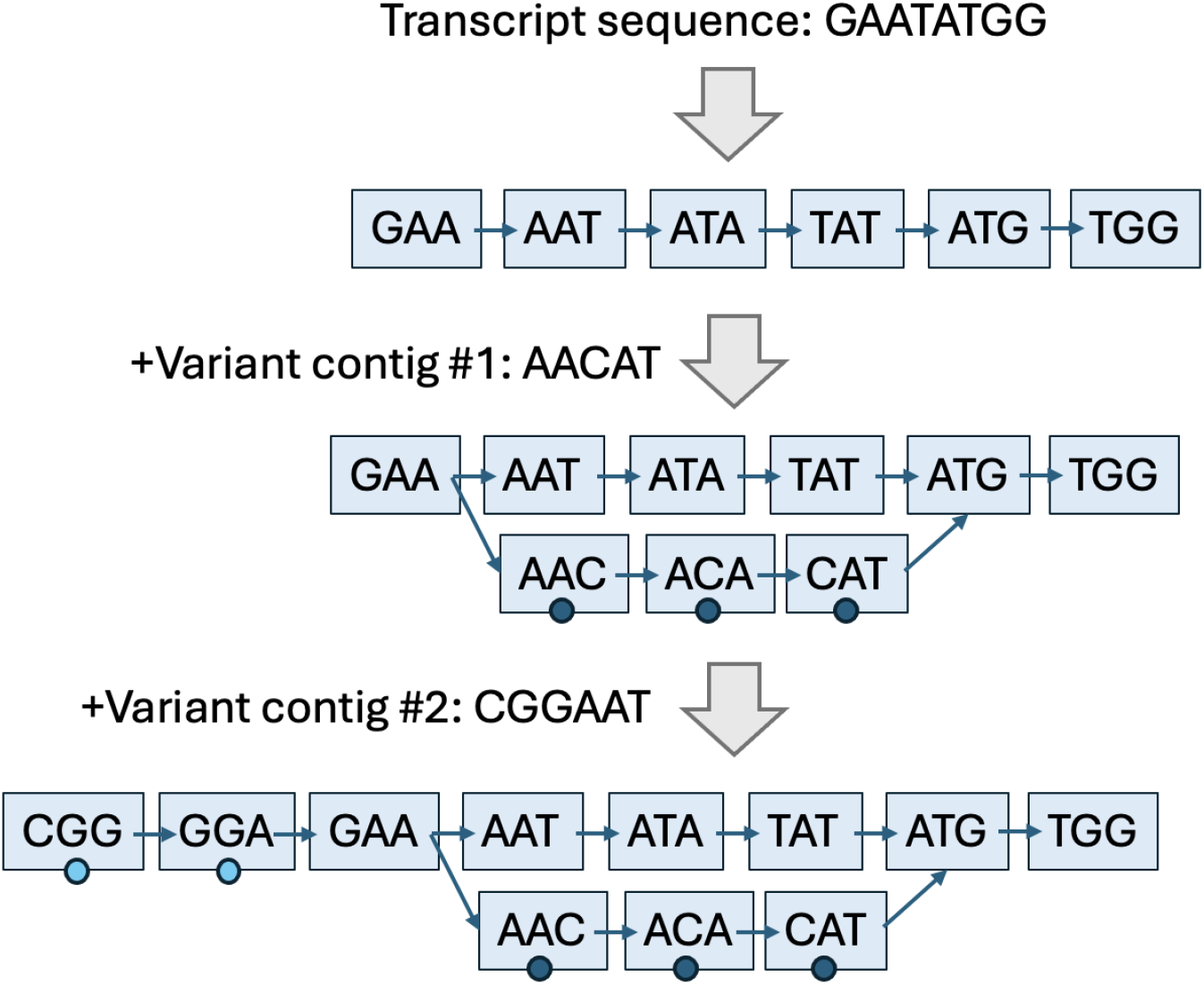
Building a de Bruijn graph with shades. One transcript sequence (parent color) and two variant contigs (shades, shown as circles) are represented. In the end, 11 k-mers (k=3) are present, all with the same parent color, two with one shade of that color, and three with another shade of that color.

Next, we provide a formal definition of a shade:

Let:

- *T* = {*T*_1_, …, *T*_*n*_} be the set of annotated transcripts
- [*n*] = {1, 2, …, *n*} be the set of *canonical* colors (like those used in standard pseudoalignment).
- *S* be a disjoint set of *shade* colors.
- π : *S*⟶[*n*] be a parent function, assigning each shade s∈*S* exactly one canonical parent color π(s).
- *A:=*[*n*]∪*S* be the universe of all colors.

A colored de Bruijn graph of order *k* is a directed graph *G* = (*V, E, C*) with:

- Each vertex *ν*∈*V* representing a unique *k*-mer over the alpha Σ = {*A, T, C, G*}.
- A directed edge (*ν, w*)∈*E* whenever the (*k* − 1)-suffix of *ν*’s *k*-mer equals the
- (*k* − 1)-prefix of *w*’s *k*-mer.
- *C*(*ν*)⊆*A* as the color set of each vertex *ν*.

We call s∈*S* a shade based on the following:

- Parent relationship: Each shade *s* is associated with exactly one canonical transcript color via a parent mapping π(*s*) ∈ [*n*]; that is, *s* has exactly one parent color.
- Parent inclusion: A shade never appears without its parent color on any *k*-mer:

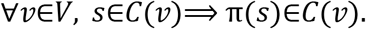

When adding a shade to a transcript de Bruijn graph, the shade is added if and only if the *k*-mer does not exist in the original graph with its parent color. Note: Each shade implicitly adds its parent color to the *k*-mer color set according to the following algorithm:

#### Algorithm

Add a new sequence to the colored de Bruijn graph as a shade

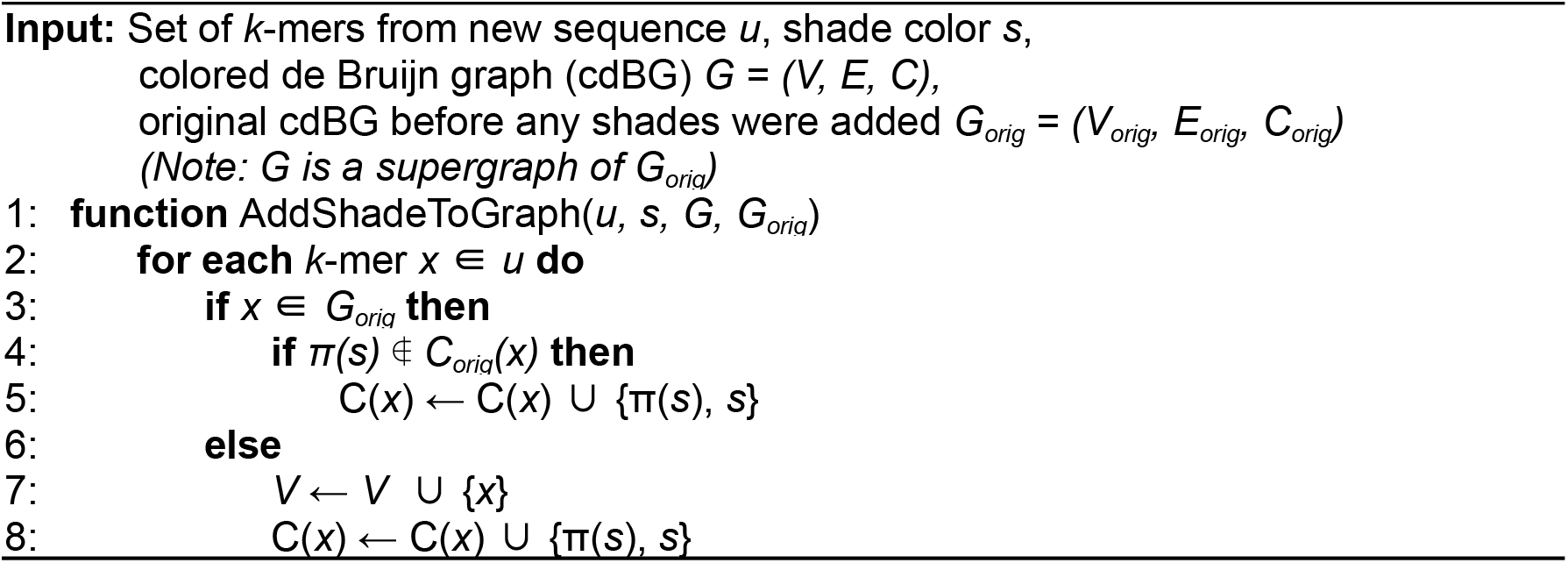

To create equivalence classes, first set-intersection is done among the parent colors encountered in the read then set-union is performed among the shade colors corresponding to those parent colors. See the following algorithm.

#### Algorithm

Assign equivalence class to read

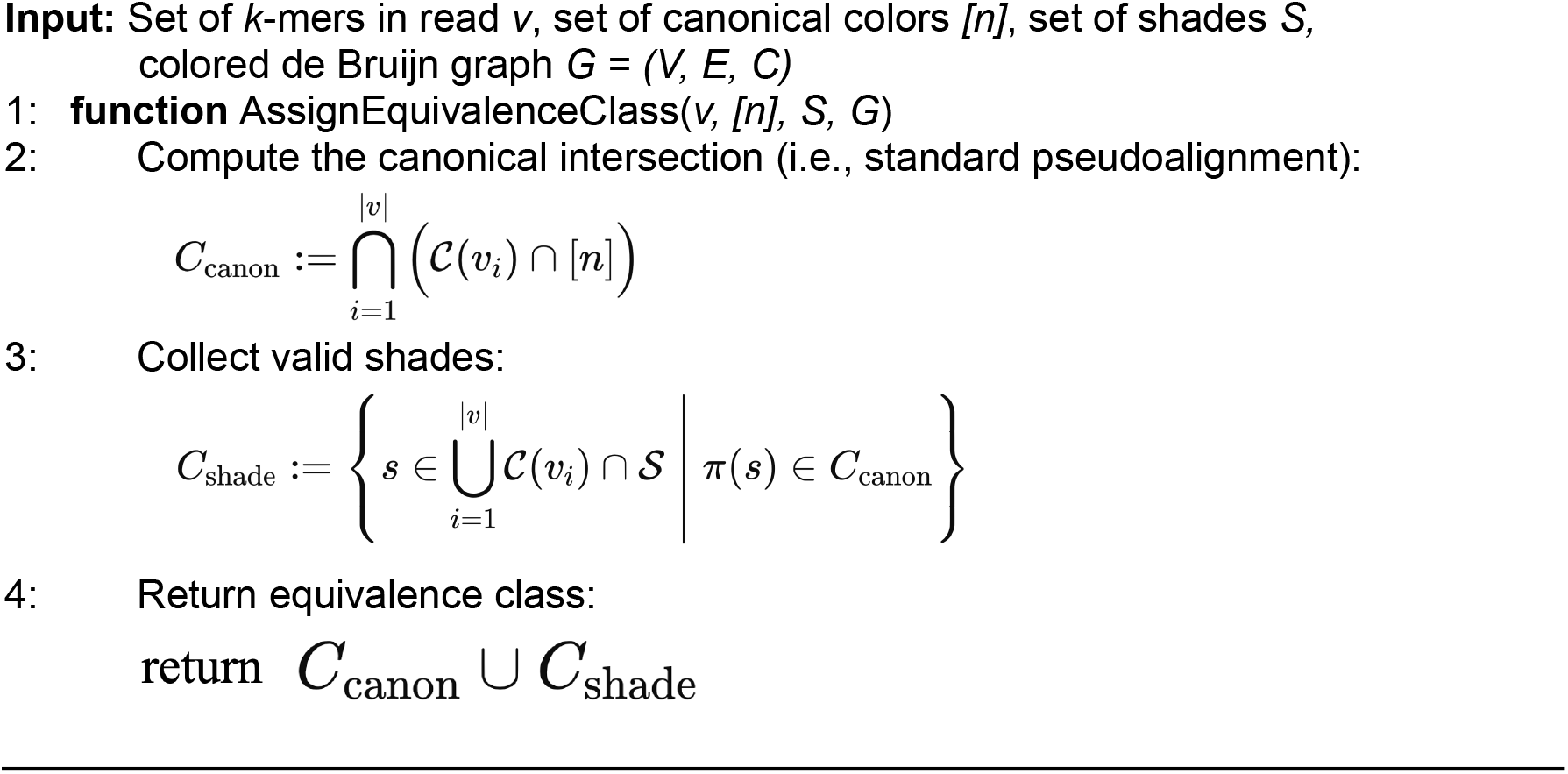

### A general-purpose k-mer toolkit

klue is a general-purpose k-mer toolkit that internally uses a colored compacted de Bruijn graph (ccdBG), implemented via the C++ Bifrost library (Holley and Melsted 2020). Each input file is assigned a unique color, encoding sample identity in the graph. A given k-mer may appear in multiple samples and thus exhibit a multi-color profile. The ccdBG compacts adjacent k-mers into longer contiguous sequences called unitigs, conserving memory and enabling analysis over longer sequence contexts.

A core feature of klue is its ability to extract contigs—unitigs, or contiguous substrings of unitigs, that share the same color profile. For instance, given three input samples colored red, blue, and green, klue can extract red-only contigs, red+blue shared contigs, contigs found in any two colors, and so on. A comprehensive suite of set operations over input samples is supported by klue. Figure 2 shows an example of extracting contigs based on color profile, wherein monochromatic contigs (those unique to one sample) are extracted. These contigs can represent tumor-specific mutations, species-specific sequences, or treatment-induced transcripts. The resulting contigs are written to FASTA files, where they can be subjected to further downstream analyses such as mapping, annotation, or pseudoassembly.

**Figure 2:**
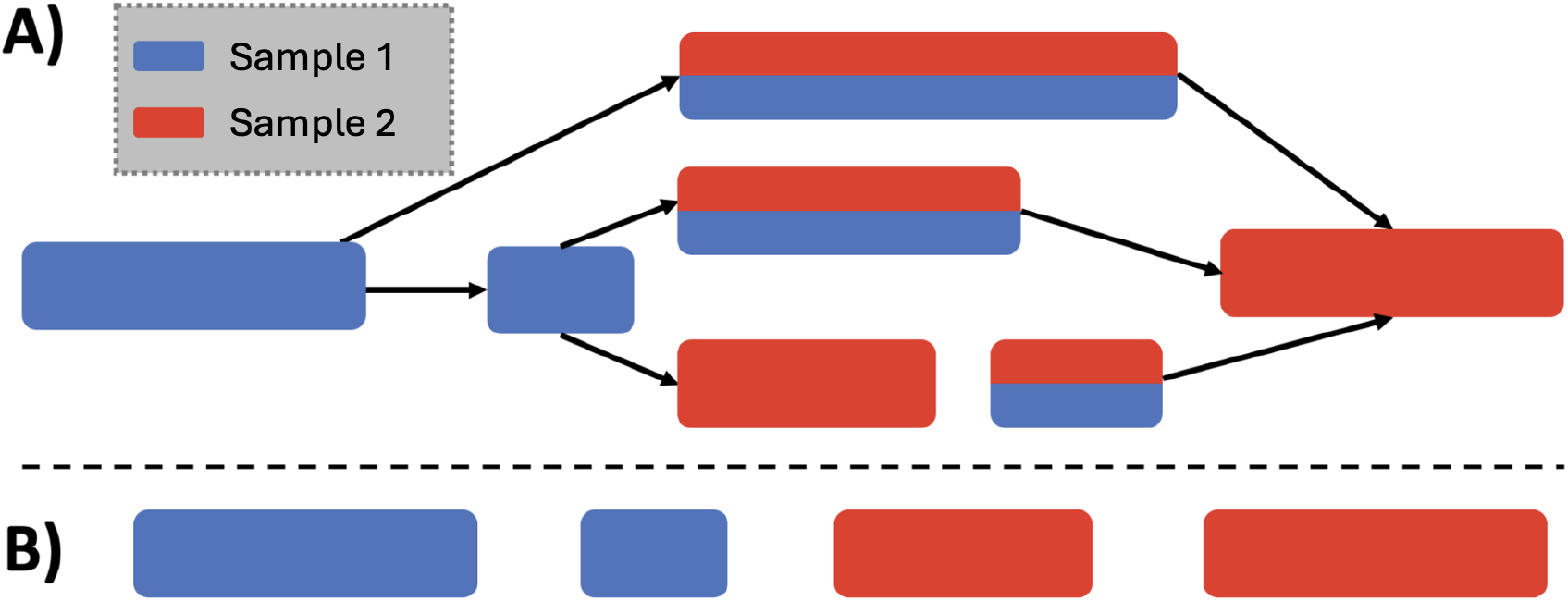
Partitioning a colored compacted de Bruijn graph. (A) The structure of the graph. The boxes shown are the nodes of the graph and represent colored contigs. (B) Monochromatic contigs that are extracted.

To illustrate klue in practice, we applied the method to eight different mouse genome assemblies (Ferraj et al. 2023), including the standard C57BL/6J mouse reference genome assembly (GRCm39). As expected, the genomes of the five inbred laboratory mouse strains (C57BL/6J, A/J, 129S1/SvImJ, NOD/ShiLtJ, NZO/HlLtJ) are more similar to each other than to the genomes of the three inbred wild-derived strains (WSB/EiJ, PWK/PhJ, CAST/EiJ) based on overlap of k-mer content from the klue-extracted k-mers (Figure 3), recapitulating expected phylogeny (Morgan and Welsh 2015). This shows that in addition to pseudoassembly, klue can be used as a convenient tool for extracting shared and unique k-mers.

**Figure 3:**
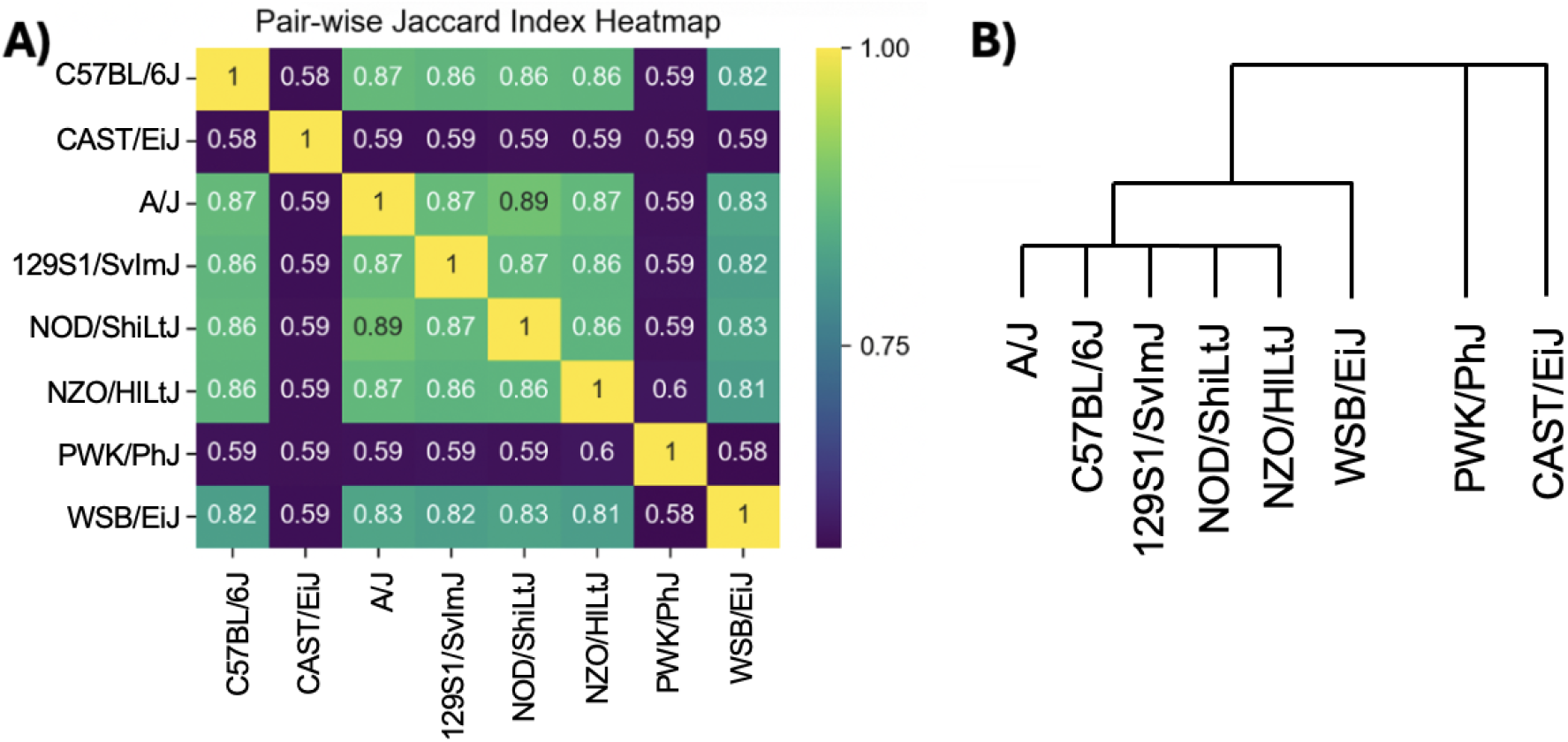
Relationship between different mouse strains. (A) Jaccard similarity index of k-mers between different mouse strains as determined by klue: the cardinality of the set intersection, X, represents the number of k-mers shared between two mouse strains while the cardinality of the set union, Y, represents all the number of k-mers present in either or both of two mouse strains; the Jaccard index is the ratio of X to Y. (B) The phylogenetic relationship between the 8 mouse strains.

Next, we evaluated klue’s ability to infer cell identity using data from single-nucleus RNA-seq experiments conducted on multiple tissues from multiple mouse strains (Rebboah et al. 2025) as part of work of the IGVF Consortium (IGVF Consortium 2024). Specifically, we examined kidney tissue from eight A/J mice and eight PWK/PhJ mice. Contigs unique to each mouse strain genome assembly were extracted, and only 61-bp contigs were retained, as these correspond to single nucleotide polymorphisms (SNP)s when using a k-mer size of 31. We focused on k-mers overlapping SNPs rather than all strain-specific k-mers because these “SNP contigs” included both the variants and their conserved flanking regions. These common anchors helped reduce noise from spurious k-mers that can arise due to artifacts in genome assemblies or differences in sequencing quality between strains. We then used kallisto (Melsted et al. 2021; Sullivan et al. 2023) to map reads to the strain-specific contigs, and cell-level strain assignments were determined based on the number of UMIs mapping to each strain (Figure 4). This klue+kallisto demultiplexing method could successfully resolve cells from PWK/PhJ mice versus A/J mice (Figure 5).

**Figure 4:**
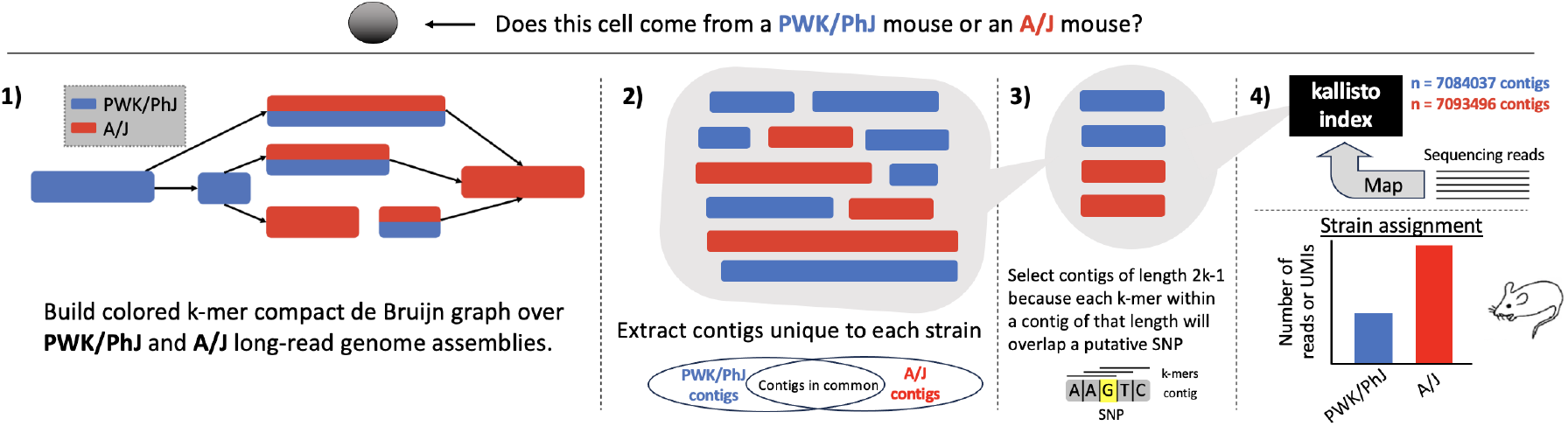
klue for mouse strain demultiplexing. Overview of the workflow used for strain demultiplexing with klue and kallisto.

**Figure 5:**
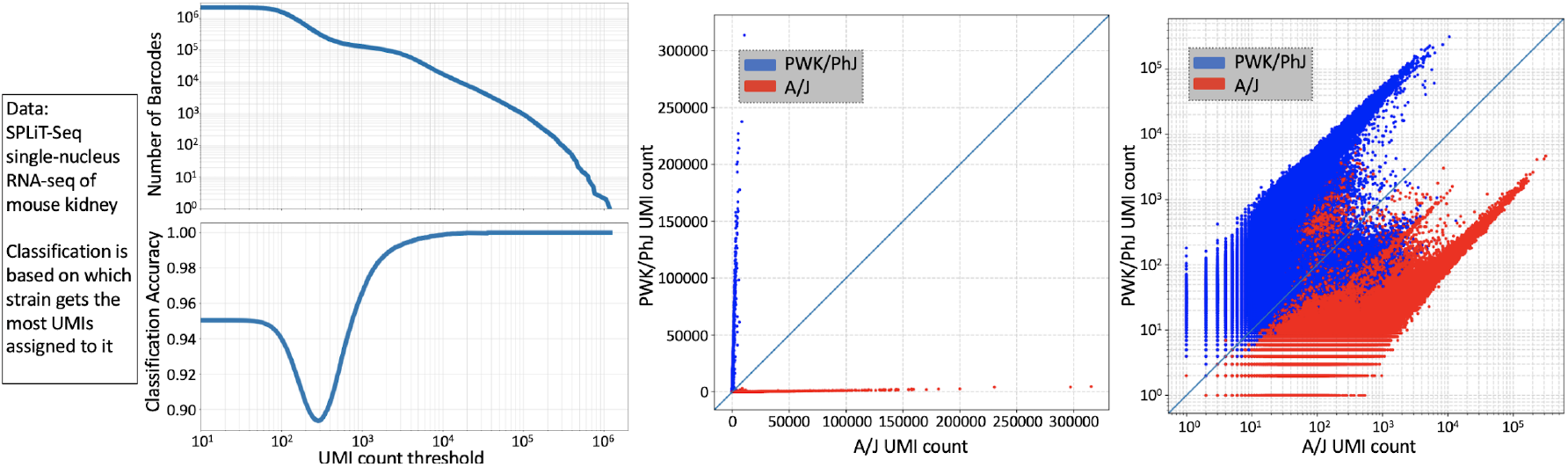
Results of klue+kallisto demultiplexing of the PWK/PhJ and A/J mouse cells. Accuracy could be determined because the PWK/PhJ cells and A/J mouse cells were placed into separate wells at the initial step of the split-pool barcoding, therefore the well barcode could serve as a ground truth label for each cell.

**Figure 5a:**
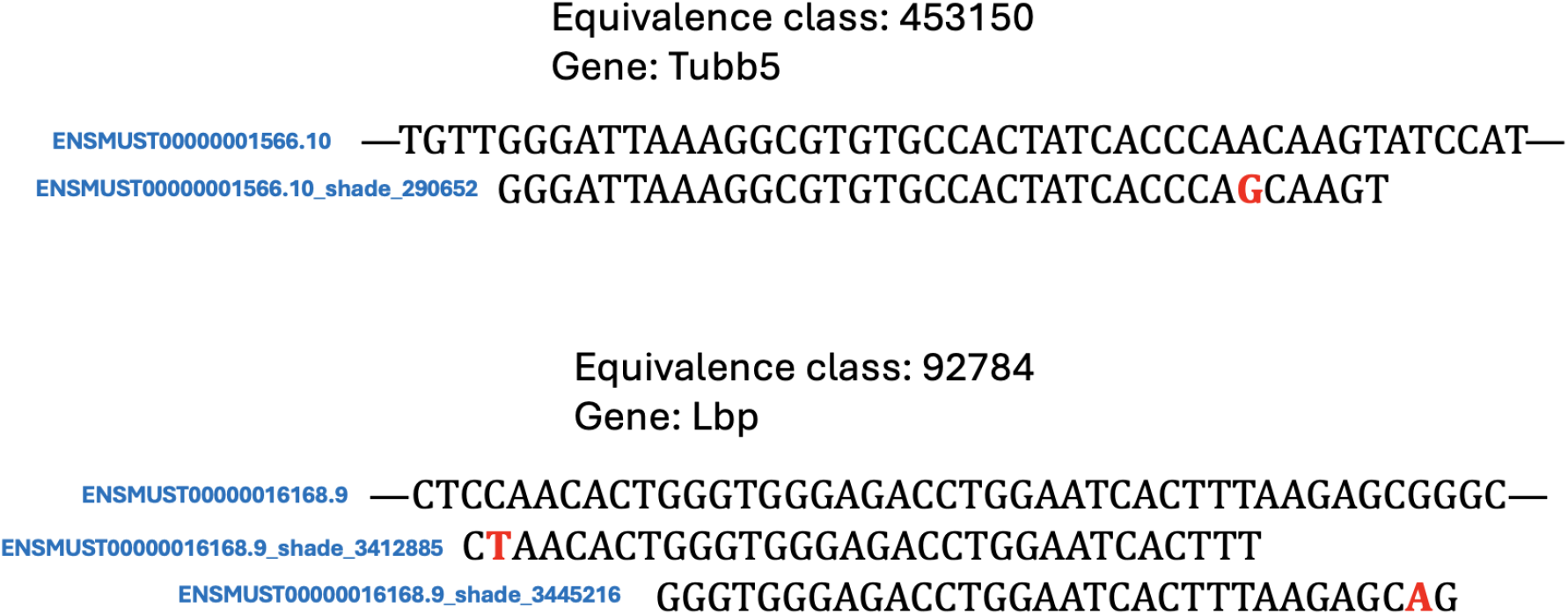
Examples of equivalence classes containing shades. The equivalence classes corresponding to two different genes are shown, each containing a standard transcriptome target and one or more “shade” targets.

### Application to cancer genomics

#### A mouse melanoma model

We performed pseudoassembly on single-cell data from a mouse melanoma model in order to identify cell-type specific mutations directly from sequencing reads. The dataset we used (Sun et al. 2019) featured 10x Genomics (version 2 chemistry) single-cell RNA-seq of mouse melanocyte stem cells (McSCs) and melanomas arising from the McSCs. In the (Sun et al. 2019) study, McSCs, derived from transgenic Tyr-CreER:Braf:Pten:Tomato mice, were transplanted into immunodeficient nude mice, and tumorigenesis was achieved by tamoxifen induction. Control McSCs were obtained from the telogen back skin of Tyr-CreER:R26R-Tomato mice. Single-cell RNA-seq was performed on both the tumors (melanomas) and the control McSCs, with three biological replicates per condition; replicates were pooled and not distinguished in downstream analyses in order to increase the ability to detect mutations. Single-end bulk RNA-seq of control McSCs was also performed.

To identify cell-type specific mutations, klue was applied to four files:

1. A FASTA file containing both the human genome FASTA (from the T2T-CHM13v2.0 assembly) and a transcriptome FASTA derived from the GRCh38 assembly.
2. Bulk RNA-seq FASTQ file for control McSCs.
3. Single-cell RNA-seq FASTQ file for control McSCs.
4. Single-cell RNA-seq FASTA file for McSC-derived melanomas.

The reason for including all these datasets was to identify melanoma-unique contigs (containing 31-mers unique to file #4), that would not include barcodes, adapters, or other sequences common to 10x experiments (removed by inclusion of dataset #3), any existing sequences in the genome (removed by inclusion of dataset #1), or sequences from exon-exon junctions (removed by inclusion of datasets #2 and #3).. The resultant contigs were mapped to the human transcriptome using kallisto (with a k-mer size of 29) to assign “shades” (Figure 5). Finally, the melanoma single-cell RNA-seq reads were mapped (using kallisto version 0.51.1) to an index containing the human transcriptome targets along with their shades.

To obtain cell clusters, the melanoma single-cell RNA-seq reads were also mapped to a standard kallisto index of the human transcriptome. After quantification of the data with kallisto | bustools (Melsted et al., 2021; Sullivan et al., 2024) to generate a cell-by-gene count matrix, we filtered for cells with at least 5000 UMIs. The counts were then normalized with CP10k (Booeshaghi et al. 2022) normalization followed by the Scanpy (Wolf, Angerer, and Theis 2018) default log1p transformation (Booeshaghi and Pachter 2021). Highly variable genes were identified using the Scanpy default (Rich et al. 2024) and then nearest neighbor graphs were constructed from the cell coordinates on the top 40 principal component analysis (PCA) subspace using Scanpy. The Leiden algorithm (Traag, Waltman, and van Eck 2019) was applied using Scanpy, resulting in 15 clusters. Clusters with fewer than 50 cells were excluded, resulting in a final count of 2776 cells distributed across 10 clusters. As the number of clusters obtained was larger than that obtained in the original study which produced the dataset (Sun et al. 2019), we merged related clusters to create more coarse-grained groupings. The original study performed pseudotemporal trajectory analysis (Trapnell et al. 2014) to define the branching transition between McSC cells to either a mesenchymal-like cell type or a neural crest / neuronal-like cell type. We therefore merged two clusters with high expression of mesenchymal markers (Dcn, Col1a1, Col1a2), merged five clusters with high expression of neural crest/neuronal genes (Nes, Foxd3, L1cam, Ngfr), and then merged the remaining three clusters which had high expression of the genes Fosb, Klf4, Serpine2, Cdkn1a from a published metastatic melanoma gene set (Perego et al. 2018). These correspond to the “mesenchymal-like” cluster, the “neural crest / neuronal-like” cluster, and the “intermediate” cluster, respectively, from the original publication (Figure 6). The original study also identified two additional clusters: a neural crest / neuronal-like cluster characterized by high expression of proliferation genes (Mki67, Cdk1), which we merged into the broader neural crest / neuronal-like cluster, and a cluster composed of control McSC cells. However, since we did not include control McSC data in our cell type clustering analysis, and the few McSC cells present in the melanoma dataset were likely removed when filtering out clusters with low cell counts, this cluster was not represented in our analysis.

**Figure 6:**
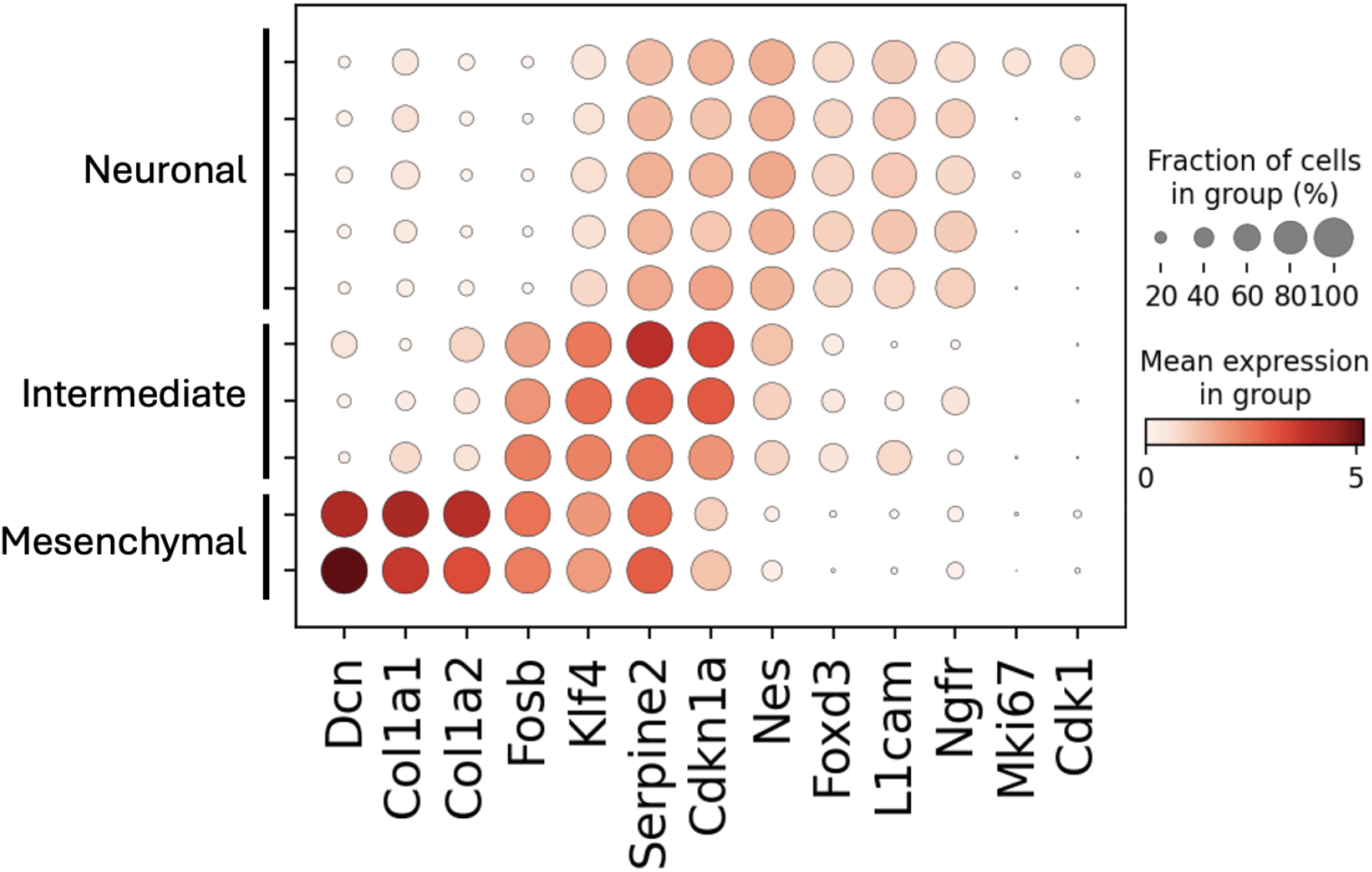
Cell type clustering of melanoma single-cell RNA-seq. Based on expression of marker genes, three coarse-grained clusters of cells with a mesenchymal-like signature, cells with a neural crest / neuronal-like signature, and intermediate cells with a signature between neuronal and mesenchymal could be resolved.

Next, we further analyzed the derived clusters, defined by gene-level counts, using the transcript compatibility counts (TCCs) from read mapping to the index containing the shades. We sought to identify mutations that were expressed uniquely in certain clusters, i.e., mutation cell-type specificity. We first filtered the cell-by-TCC count matrix to only contain equivalence classes (ECs) which contained a shade target mapping and which were present in at least 10 cells. All ECs corresponding to multiple genes, unannotated genes, or pseudogenes were excluded. For differential expression testing of shade counts, a *2 × 2* contingency table for each shade-containing EC was constructed as follows:

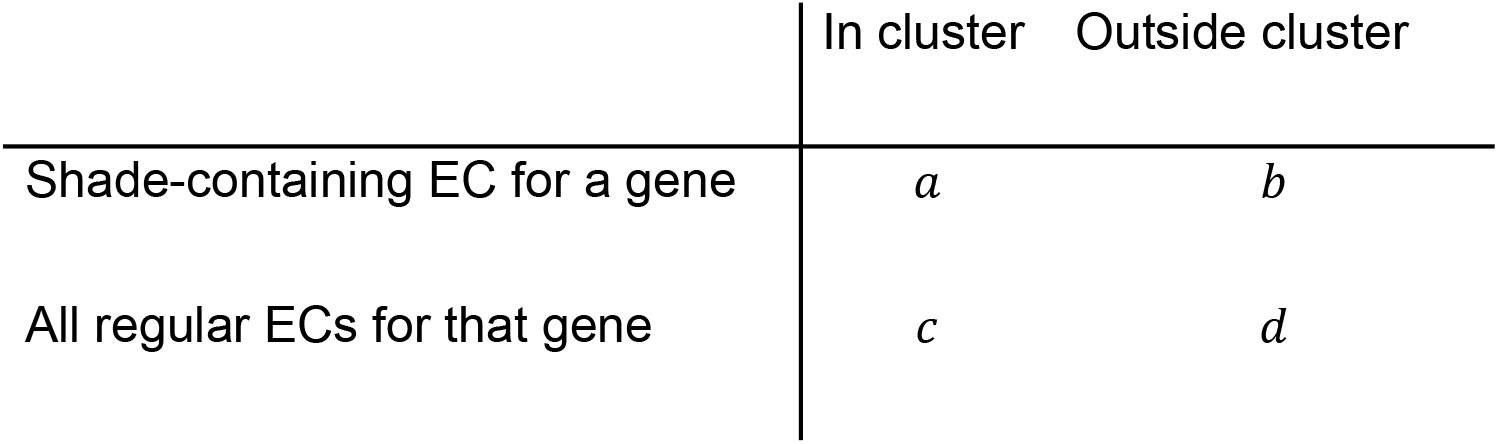

|  | In cluster | Outside cluster |
| --- | --- | --- |
| Shade-containing EC for a gene | $a$ | $b$ |
| All regular ECs for that gene | $c$ | $d$ |

Here, *a, b, c*, and *d* represent cell numbers, i.e., the number of cells that contain at least one count for the EC. The odds ratio is then calculated as follows, with higher values indicating greater specificity of the shade (i.e., the putative mutation) for the cluster being evaluated:

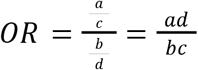

For each EC a *p*-value was determined using Fisher’s exact test. A volcano plot depicting the results of differential expression testing of shade (mutation) counts is shown in Figure 7.

**Figure 7:**
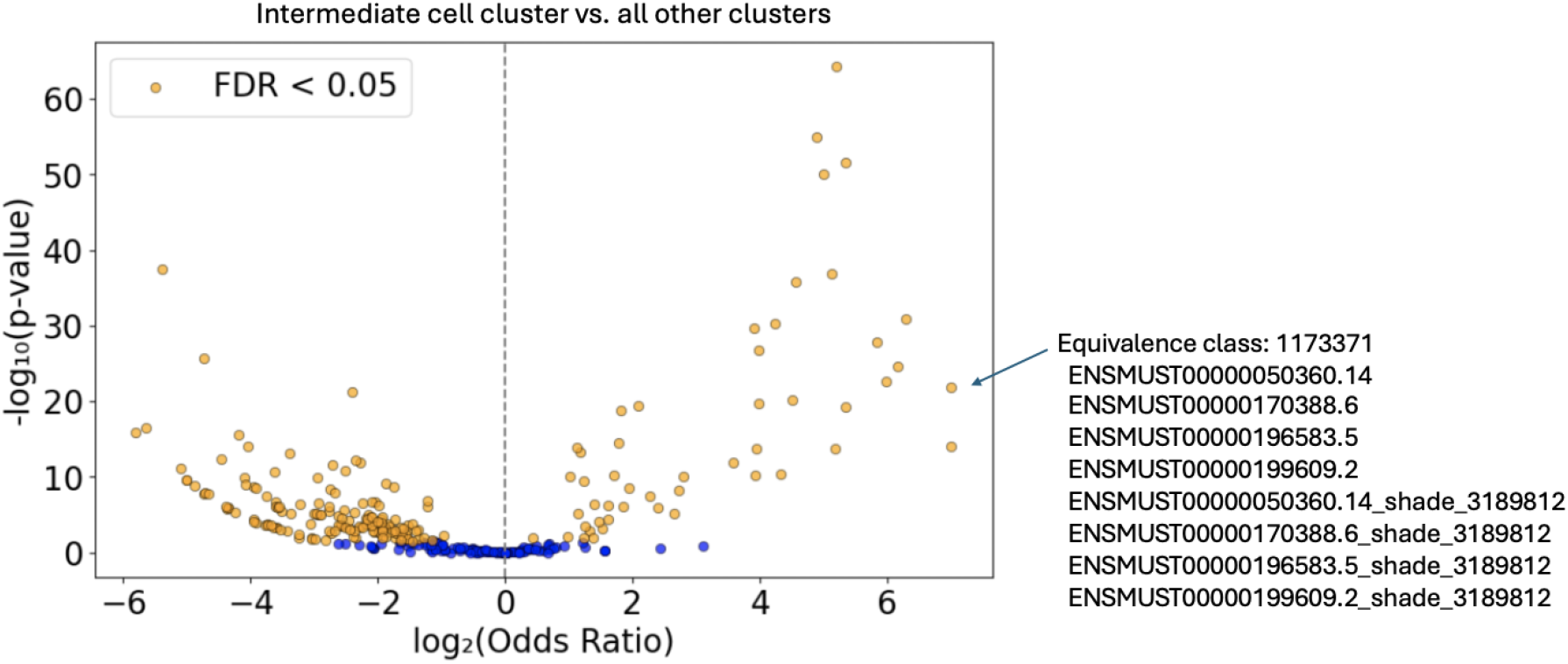
Differential expression testing of shade (mutation) counts. Each point on the volcano plot represents a shade-containing equivalence class (EC). An example of such an EC (EC number 1173371), corresponding to the P2ry12 gene, is shown on the right. 50 ECs had an odds ratio greater than 1 and a Benjamini-Hochberg FDR-adjusted p-value less than 0.05, while 278 ECs had an odds ratio less than 1 with an adjusted p-value below the same threshold.

The method was able to identify many examples in which a shade for a given gene is abundant in one melanoma cluster but not the others (Figure 8). Specifically, the P2ry12 gene appeared to contain a differential variant in the intermediate cluster, and the Tubb5 gene was revealed to contain a differential variant in the mesenchymal cluster. These observations would not have been possible within the standard single-cell RNA-seq analysis framework (Chen et al. 2016).

**Figure 8:**
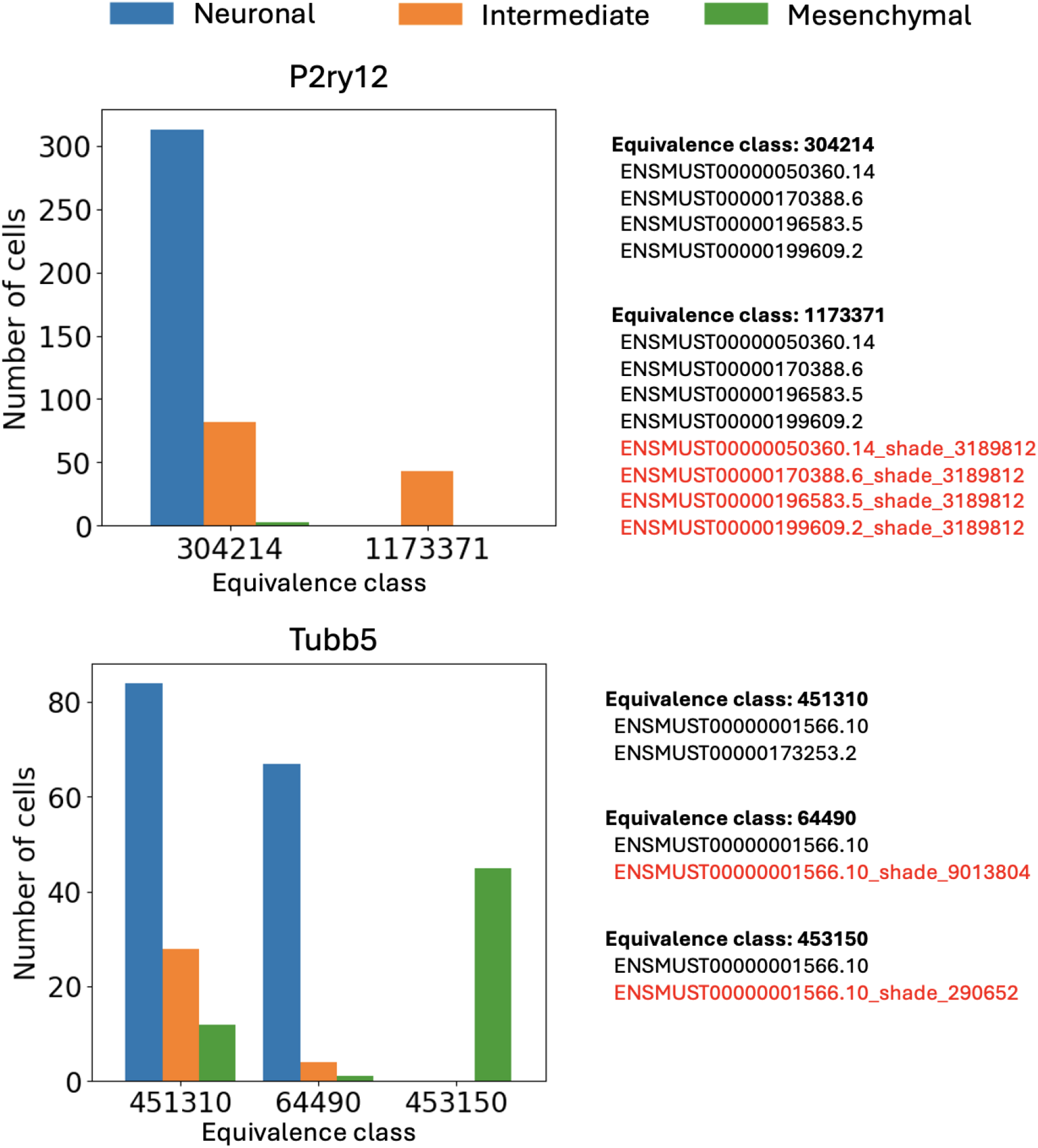
Examples of genes where the associated equivalence classes display different patterns of cluster-specific expression. Shaded targets are marked in red. A P2ry12 mutation is found specifically in the intermediate cluster; a Tubb5 mutation is found specifically in the mesenchymal cluster. The y-axis represents the number of cells for which at least one UMI is identified as being associated with the given equivalence class.

## Discussion

The (Sun et al. 2019) melanoma dataset was well-suited for identifying mutation cell type specificity due to two key factors: the inclusion of a control sample and the nature of 10x Genomics chemistry, which sequences only the ends of transcripts. The control sample was essential to distinguish true mutations from common genetic variants such as SNPs, which would otherwise be picked up as shades. While sequencing only the 3’ ends of transcripts limits coverage and may miss mutations located in the middle of genes, it also helps reduce false signals. For instance, incomplete coverage might result in certain transcript regions being captured only in the disease group but not in the control group—leading to spurious disease-unique contigs. Taken together, the (Sun et al. 2019) datasets provide a compelling argument for pseudoassembly to discover cell-type specific mutations. The generality and flexibility of klue makes it suitable for any (single-cell) genomics datasets, and it should prove to be a useful complement to standard assembly algorithms.

## Data and code availability

The data and code needed to reproduce the results in this paper are available at: https://github.com/pachterlab/SBP_2025

The klue program is available at https://github.com/pachterlab/klue

## Author Contributions

The pseudoassembly idea was conceived by DKS. DKS implemented klue with the assistance of MB. Results were obtained by DKS. The manuscript was drafted by DKS, and edited by DKS and LP.

## Acknowledgements

We thank Ali Mortazavi, Elisabeth Rebboah, Fairlie Reese, and Ryan Weber for help with testing klue. DKS was funded in part by NIH National Institute of General Medical Sciences (UCLA–Caltech Medical Scientist Training Program T32 GM152342). MB and LP were funded in part by NIH UM1 HG012077.

